# Secondary bile acid production by gut bacteria promotes Western diet-associated colorectal cancer

**DOI:** 10.1101/2023.03.17.533140

**Authors:** Esther Wortmann, Annika Osswald, David Wylensek, Stephanie Kuhls, Olivia I. Coleman, Quinten Ducarmon, Wei Liang, Nicole Treichel, Fabian Schumacher, Colin Volet, Silke Matysik, Karin Kleigrewe, Michael Gigl, Sascha Rohn, Burkhard Kleuser, Gerhard Liebisch, Angelika Schnieke, Rizlan Bernier-Latmani, Georg Zeller, Dirk Haller, Krzysztof Flisikowski, Soeren Ocvirk, Thomas Clavel

**Author notes:** shared first authors (Esther Wortmann, Annika Osswald). shared last authors (Soeren Ocvirk, Thomas Clavel). **Conflict of Interest** The authors declare no potential conflicts of interest.

## Abstract

Western diet is an important risk factor for the development of sporadic colorectal cancer (CRC). Dietary fat stimulates bile acid (BA) production by the host and their conversion to secondary BAs by 7α-dehydroxylating (7αDH+) bacteria, but causal proof of their tumor-promoting effects *in vivo* is lacking. To address this, we performed feeding studies in a genetically engineered pig model of CRC combined with multi-omics analyses and gnotobiotic mouse studies. Western diet worsened the disease phenotype in *APC*^1311/+^ pigs. This was accompanied by microbiota changes, increased levels of the secondary bile acid deoxycholic acid (DCA), and higher colonic epithelial cell proliferation. The latter was counteracted by using the BA-scavenging drug colestyramine. Metagenomic analysis across multiple cohorts revealed higher occurrence of *bai* (BA inducible) operons from *Clostridium scindens* and close relatives in stool of CRC subjects (n = 1,034). Using two gnotobiotic mouse models of CRC, we demonstrate that colonization with 7αDH+ bacteria (*C. scindens* or *Extibacter muris*) increased colonic tumor loads. This work provides clear evidence for the causal role of microbiome-derived DCA production in CRC under detrimental dietary conditions, opening avenues for future preventive strategies.

## Introduction

Colorectal cancer (CRC) has one of the highest incidences and mortalities among all cancer types worldwide, with approximately 1.9 million new cases and 0.9 million deaths in 2020 (Sung et al., 2021). Besides genetic factors, including mutations in the tumor-suppressing adenomatous polyposis coli (*APC*) gene, sporadic CRC risk is predominantly influenced by environmental and lifestyle factors, including diet and the gut microbiome (Huxley et al., 2009; Keum & Giovannucci, 2019; Lichtenstein et al., 2000). There is convincing evidence for a positive association between CRC risk and Western-style diets rich in fat and red/processed meat, whilst high-fiber intake is negatively associated with CRC risk (Chen et al., 2013; Garcia-Larsen et al., 2019; Reynolds et al., 2019). In healthy individuals in South Africa, who have a very low CRC risk, a diet switch every two weeks from a traditional high-fiber/low-fat to a Western-style low-fiber/high-fat dietary pattern altered gut microbiota composition and metabolism, leading to an increase in markers associated with CRC risk such as intestinal epithelial cell (IEC) proliferation and infiltration of CD3+ and CD68+ immune cells (O’Keefe et al., 2015). Alaskan Native people, who traditionally consume a diet rich in fat and have the world’s highest risk for CRC, have a gut microbiota characterized by lower diversity and metabolic adaption to bile acid (BA) metabolism (Ocvirk et al., 2020).

High fat intake stimulates BA synthesis by the host, resulting in increased amounts of BAs entering the colon, where they are converted to secondary BAs by gut bacteria. Elevated levels of secondary BAs, in particular deoxycholic acid (DCA), were detected in feces of CRC patients and healthy cohorts at high CRC risk, and they correlated with higher occurrence of BA-inducible (*bai*) genes involved in BA 7α-dehydroxylation (7αDH) by gut bacteria (Hill et al., 1975; Kawano et al., 2010; Ocvirk & O’Keefe, 2021; Ou et al., 2013; Wirbel et al., 2019). In the murine *Apc*^min/+^ model, a diet supplemented with the primary BA cholic acid (CA, 0.4% w/w, 12 weeks), the precursor of DCA, increased the number and size of intestinal tumors (Wang et al., 2019). This effect was prevented by antibiotic treatment, suggesting a contribution of the gut microbiota to CA-associated tumorigenesis in this CRC model. Oral intake of DCA promoted tumor development in the proximal colon of wildtype mice (0.2% w/w in diet for up to 10 months) (Bernstein et al., 2011), high-grade dysplasia in AKR/J mice treated with azoxymethane (AOM) (0.25% w/w in diet for >10 weeks) (Flynn et al., 2007) and higher numbers of intestinal tumors in *Apc*^min/+^ mice (0.2% w/v in drinking water for 12 weeks) (Cao et al., 2014). Fecal microbiota transplantation from DCA-treated *Apc*^min/+^ donor mice to microbiota-depleted *Apc*^min/+^ recipients on a control diet induced increased intestinal tumor numbers (Cao et al., 2017). However, in these studies, mice were fed with DCA directly, which does not mirror physiological conditions, or DCA levels were not analyzed. Moreover, the *Apc*^min/+^ mouse model develops tumors primarily in the small intestine, which does not reflect the location of tumorigenesis in humans and contrasts to colon being the main site of secondary BAs production (A.R Moser et al., 1995; Amy Rapaich Moser et al., 1990). Hence, clear evidence for a causal role of microbially produced DCA in relation to diet and underlying mechanisms in colonic tumorigenesis is lacking.

Here we investigated the role of secondary BA production by gut bacteria in CRC, based on a multi-layer approach using dietary interventions in genetically engineered pigs, patient stool samples, gnotobiotic mouse models of CRC, and a combination of multi-omics and cultivation methods. This allowed us to span analyses from microbial community level to detailed investigations of specific microbiota members involved in the conversion of BAs, collectively demonstrating a diet-driven, tumor-promoting function of secondary BA-producing bacteria.

## Results

### Western diet promotes high levels of secondary BAs and aggravates disease in *APC*^1311/+^ pigs

Pigs and humans are omnivores and share similarities in their gut microbiota, gastrointestinal anatomy, physiology, and metabolic processes. This has made the pig an ideal model to study gastrointestinal diseases (Gonzalez et al., 2015; Rose et al., 2022). In contrast to transgenic mouse models based on the *Apc* gene, *APC*^1311/+^ pigs recapitulate key features of human disease, including the location of tumors in the large intestine (Flisikowska et al., 2012), disease progression, and differences in severity within families (Flisikowski et al., 2022).

To investigate the role of a Western diet on the development of CRC under controlled conditions in a preclinical setting, we performed a feeding intervention in *APC*^1311/+^ pigs. The animals were fed (1 kg/d) either a control diet (CTRL) or an experimental diet enriched in red meat and lard (RL diet) for 3 months (see experimental design in **Fig. S1a** and diet composition in **Table S1**). Compared with control *APC*^1311/+^ pigs, both the number and size of colonic polyps were significantly increased in the RL diet group (**Fig. 1a**). Higher numbers of Ki67+ cells in distal colonic crypts indicated increased proliferation of the epithelium (**Fig. 1b**).

**Fig 1.**
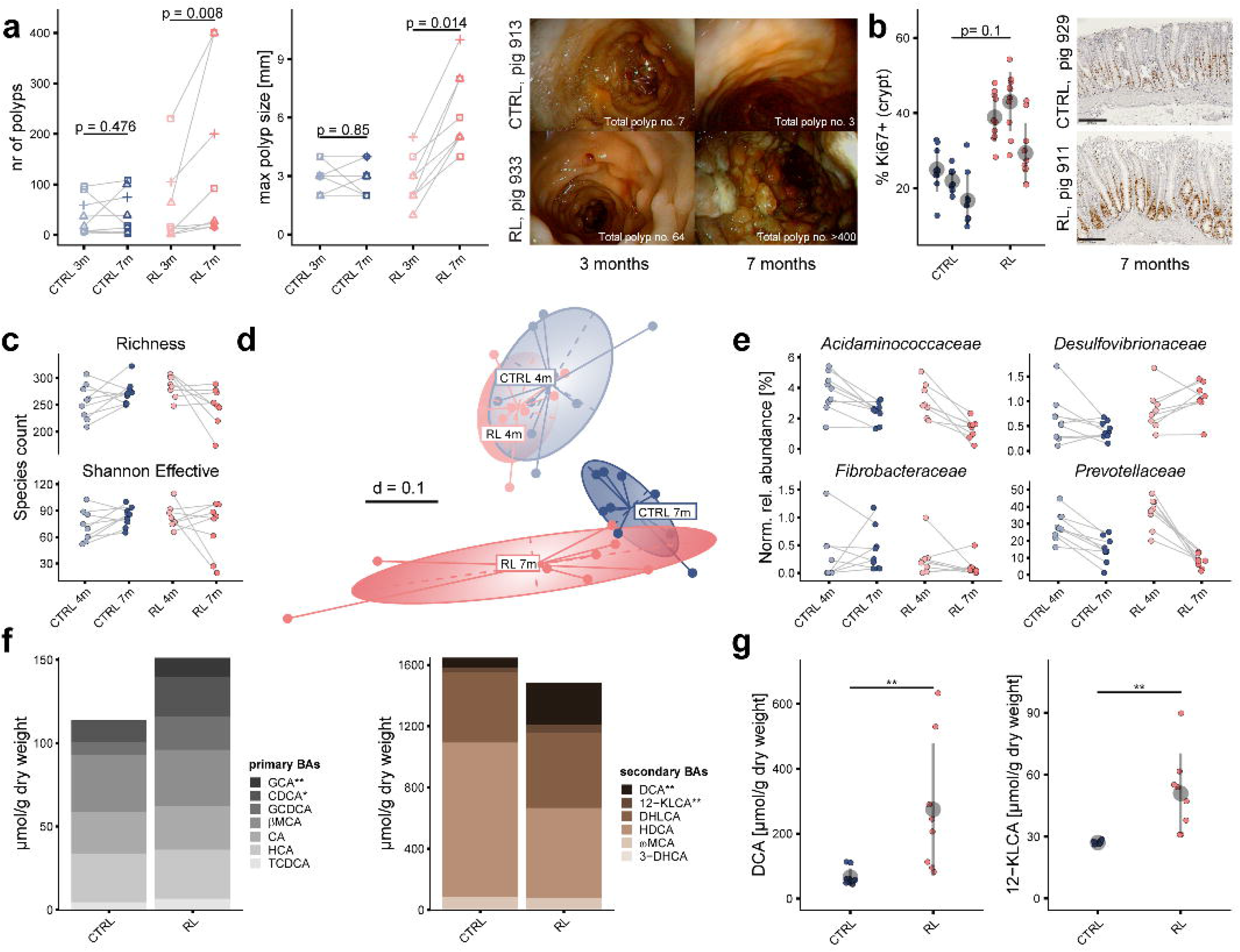
Effects of Western diet on disease phenotype, fecal microbiota (16S rRNA amplicon sequencing) and bile acids (LC-MS/MS): **a,** number and size of polyps in the distal colon of *APC*^1311/+^ pigs aged 3 months (3m, 1 month prior to the feeding intervention) or 7 months (7m, end of feeding intervention) and fed either the red meat and lard diet (RL, n = 8; red symbols) or the control diet (CTRL, n = 9; blue). Litters are indicated by different symbols. Right panel: Exemplary endoscopic pictures (last 40 cm of the colon). **b,** Percentage of Ki67+ cells in crypts of the distal colon from CTRL and RL pigs (n = 3 each) after the feeding intervention; for each pig, 10 randomly selected fields were counted (small colored dots). Mean values and standard deviations per pig are shown in grey. Right panel: Exemplary Ki67 staining of distal colon samples. Black bars on the pictures represent 200 µm. **c,** Richness and Shannon effective counts over time; p = 0.0206 and 0.1390, respectively (comparison of delta values). 4m (4 months) is the start of the dietary intervention, 7m (7 months) the end. **d,** *Beta*-diversity shown as metaNMDS plot of generalized Unifrac distances (p = 0.001, PERMANOVA; pairwise comparisons with Benjamini-Hochberg adjustment: start, p.adj. = 0.399, end, p.adj. = 0.0015). **e,** Relative abundances of dominant bacterial families with significantly different changes over time (end-start/start) between feeding groups (Wilcoxon rank sum test, p ≤ 0.05); **f,** Primary (grey stack bars) and secondary (brown) BA levels in faeces at the end of the feeding period; mean values per feeding group are shown (n = 9 for CTRL, n = 8 for RL). All values are available in **Table S2**. GCA = Glycocholic acid, CDCA = chenodeoxycholic acid, GCDCA= Glycochenodeoxycholic acid, *β*MCA = *β*-muricholic acid, CA = cholic acid, HCA = hyocholic acid, TCDCA = tauro-chenodeoxycholic acid, DCA = deoxycholic acid, DHCA = dehydro-lithocholic acid, HDCA= hyo-deoxycolic acid, ωMCA = ω-muricholic acid, 12-KLCA = 12-keto-litocholic acid, 3-DHCA = 3-dehydro cholic acid. **g,** Dot plots of the secondary BAs DCA and 12-KLCA in faeces. Mean values and standard deviations across all pigs per group are shown in grey. Statistics: **a,** Paired Wilcoxon signed rank test; **b,** unpaired Wilcoxon rank sum test; **c** and **e,** Wilcoxon rank sum Benjamini-Hochberg adjustment (* p.adj. ≤ 0.05, ** p.adj. ≤ 0.01).

To assess the effects of experimental feeding on the gut microbial ecosystem, fecal samples were analyzed by 16S rRNA gene amplicon sequencing before and after intervention. *Alpha*-diversity analysis revealed a trend towards increased richness and Shannon effective counts in the CTRL group, which was not observed in the RL group, including a sharp drop in diversity in the two pigs with most severe disease progression (**Fig. 1c**). According to *beta*-diversity analysis, the microbiota structure between CTRL and RL pigs overlapped at the beginning of the feeding trial (4 months) but diverged after experimental feeding (7 months, **Fig. 1d**). At the level of dominant bacterial families, a decrease in *Prevotellaceae* and *Acidaminococcaceae* over time was observed in both groups, but it was significantly more pronounced in RL pigs (**Fig. 1e**). Whilst *Fibrobacteraceae* increased in the CTRL group, RL pigs were characterized by a significant increase in *Desulfovibrionaceae*.

Besides diet-induced effects on microbiota diversity and composition, changes in fecal metabolites were assessed. Targeted LC-MS/MS measurements showed that primary BAs were overall increased in RL pigs after dietary intervention, primarily due to glycocholic acid (GCA), which was not detected in the CTRL group (**Fig. 1f**). Total concentrations of secondary BAs were slightly higher in the CTRL group, mainly driven by hyodeoxycholic acid (HDCA), shown to suppress epithelial proliferation (M. Song et al., 2020). In contrast, levels of the CRC-associated secondary BAs DCA and 12-keto-litocholic acid (12-KLCA) were substantially higher in the RL group (**Fig. 1g**).

### The bile acid sequestrant colestyramine limits Western diet-associated epithelial cell proliferation in the colon of *APC*^1311/+^ pigs

Next, we investigated whether the increase in BAs contributed to the RL diet-induced phenotype in *APC*^1311/+^ pigs. To this end, a second feeding trial was carried out, including a third group fed the RL diet supplemented with the drug colestyramine (COL; 12g/d), a bile acid scavenger that makes BAs unavailable for bacterial transformation and enterohepatic circulation (Zarras & Vogl, 1999) (see experimental design in **Fig. S1b**). Whilst the number of polyps tended to decrease after intervention in all COL pigs (**Fig. 2a**; p = 0.062), their size varied markedly between individuals in this group (**Fig. 2b**). Moreover, the number and size of polyps in the CTRL and RL groups showed inconsistent changes over time. However, immunohistochemistry staining of Ki67+ cells confirmed the proliferative effect of the RL diet in crypts of the distal colon, which was prevented by COL (**Fig. 2c**). Targeted analysis of fecal BAs at the end of the feeding period showed higher total levels of primary and secondary BAs in RL and COL pigs (p.adj. = 0.068 and 0.066, respectively), including higher concentrations of GCA (p.adj. = 0.05), allocholic acid (AlloCA) (p.adj. = 0.07), CA, and ursocholic acid (UCA) (p.adj. = 0.05) (**Fig. 2d**). Fecal concentrations of the secondary BAs DCA and 12-KLCA increased with RL diet feeding, whilst they tended to be lower under colestyramine treatment (**Fig. 2e**). Targeted measurements of BAs were complemented by a lipidomics approach. Levels of plant sterols were elevated in fecal samples from CTRL pigs and colestyramine tended to compensate for this effect in the case of cholestanol and 5α sitostanol (**Fig. 2f**). In contrast, the levels of cholesterol and derivatives as well as total fatty acids (FA) were highest in feces of COL pigs (**Fig. 2g**). The differences in FA were mainly driven by total saturated and branched chain (iso and ante-iso) FA, whereas unsaturated and long-chain FA (22-24) showed no significant differences.

**Fig. 2.**
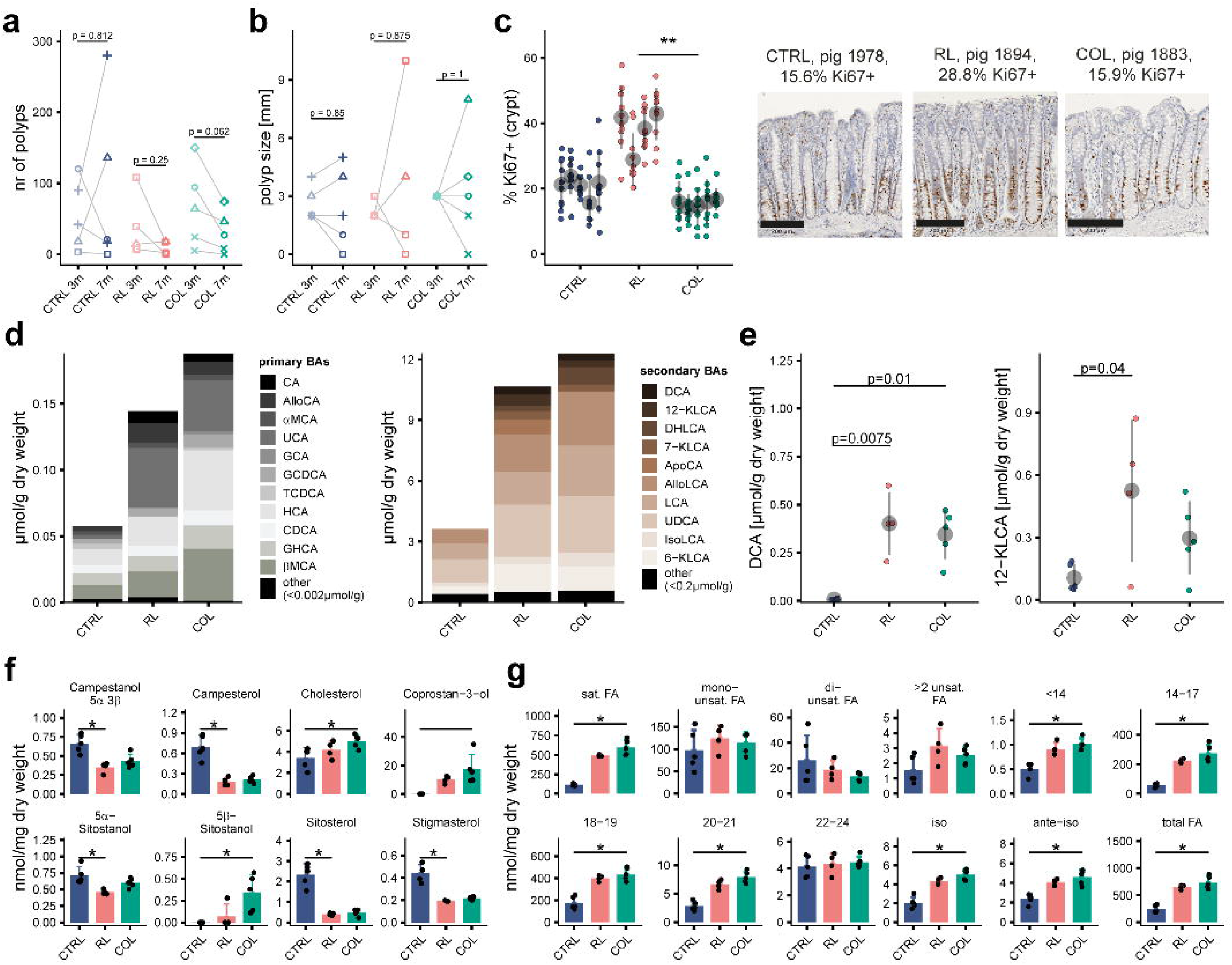
Effect of colestyramine treatment in transgenic *APC*^1311/+^ pigs: The COL diet corresponded to RL diet supplemented with 12 g/kg colestyramine (see detailed composition in **Table S1**). Polyp number (**a**) and size (**b**) in the distal colon (last 40 cm, 3m = 3 months, one month prior to intervention; 7m = 7 months, end of feeding period, n = 5 for CTRL and COL, n = 4 for RL). Litters are indicated by different symbols. **c,** Percentage of Ki67+ cells in crypts of the distal colon. For each pig, 10 randomly selected fields were counted (colored dots). Mean values and standard deviations per pig are shown in grey. Exemplary microscopic pictures of Ki67 staining (bars = 200 µm) are displayed next to the graph. **d**, Mean values of primary (grey) and secondary (brown) BAs at the end of the feeding period (7 months). All values are given in **Table S3**. **e**, Individual values for DCA and 12-KLCA. Mean values and standard deviations are show in grey. **f**, Sterol/stanol profiles. **g**, FA profiles: total saturated FA (sat. FA), monounsaturated FA, di-unsaturated FA, >2 unsaturated FA, FAs separated by chain length, branched chain FA (iso- and ante-iso), and total FA. Individual values are provided in **Table S3**. Values in **f** and **g** are shown as individual values (black dots) and mean (bars) + standard deviation (whiskers). CTRL, control diet (n = 5); RL, red meat and lard diet (n = 4); COL, colestyramine diet (n = 5). Statistics: **a** and **b**, Paired Wilcoxon signed rank test; **c-g** Kruskal-Wallis test with Dunn’s multiple comparisons and Benjamini-Hochberg adjustment (* p.adj. ≤ 0.05).

Taken together, the two feeding trials in genetically engineered pigs provide interventional evidence for the positive association between Western dietary habits, elevated levels of secondary BAs, and polyp formation. Western diet-driven epithelial cell proliferation in the colon was prevented by the drug colestyramine, which stimulated fecal excretion of lipids, making them likely unavailable for the host and BAs not accessible for bacterial conversion into secondary BAs. To further demonstrate the causal implication of secondary BAs in CRC development, we next used gnotobiotic mouse models.

### Specific 7**α**DH+ bacteria are associated with CRC

To identify the most relevant 7αDH+ bacteria in the context of CRC, we analyzed the occurrence of *bai* operons from different mOTUs (metagenomic operational taxonomic units) in fecal metagenomes from CRC patients (n = 1,034) and control individuals (n = 1,108) across multiple cohorts. The 7 mOTUs with a prevalence >2% showed specificity in terms of both, *bai* operon structure and protein sequence comparison (**Fig. 3a, b**). The occurrence of mOTUs corresponding to *Clostridium scindens* [r03437], *Clostridium hylemonae* [r11469], and the yet-unknown species *Clostridiales* sp. [m14015] was significantly higher in CRC patients (**Fig. 3c**). Cumulative relative abundances of the *bai* operons from *C. scindens* and *C. hylemonae* confirmed the relevance of these two species in CRC when analyzing all cohorts together (**Fig. 3d**). An increase was observed in nearly all individual studies, but significance depended on the cohort considered, likely due to varying size and environmental parameters (*e.g.,* diet) (**Fig. 3e**).

**Fig. 3.**
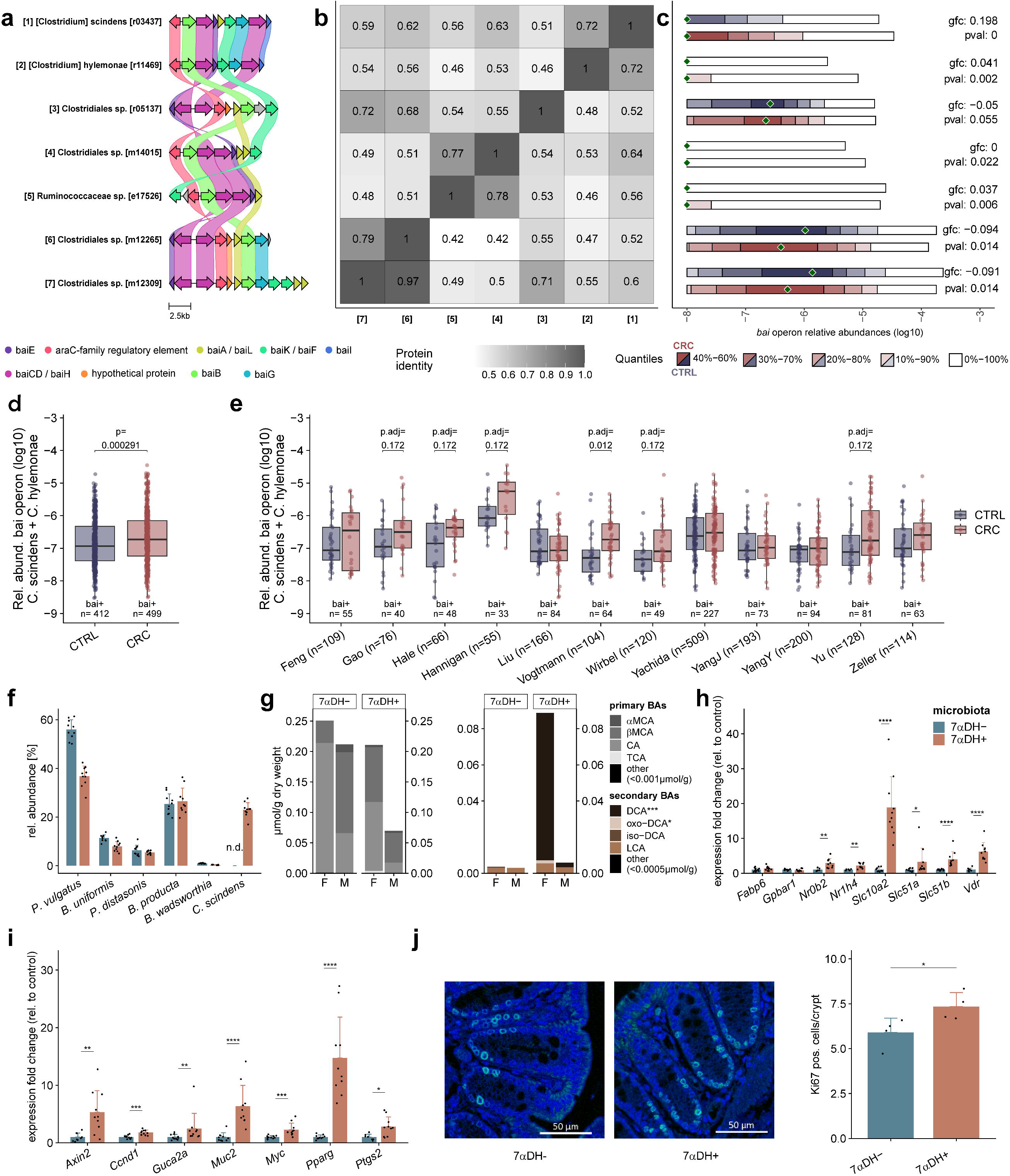
Identification of specific DCA-producing bacteria associated with CRC and functional evaluation in a gnotobiotic mouse model: **a** Genomic architecture of the *bai* operon in mOTUs (metagenomic operational taxonomic units) detected at >2% prevalence (n = 2,142). **b** Protein sequence identities between the different genomic architectures of the *bai* operon shown in **a**. **c** Quantile plots of relative abundances of mOTU-specific *bai* operons over all CRC (n = 1,034) vs. control (n = 1,108) samples. Due to high protein sequence similarity between *Clostridiales* sp. [m12265] and *Clostridiales* sp. [m12309] (last two bacteria listed), relative abundances are almost identical. **d** Cumulative relative abundance of the *bai* operons from *C. scindens* and *C. hylemonae* in samples from all cohorts. Only the samples with a detected *bai* operon from either of the two species are plotted. The corresponding number of positive samples is indicated above the x-axis. The total number of samples is indicated below the x- axis. **e** Same as in d, but per study. Studies that included <10 samples with detected *bai* operons in either of the groups (CTRL or CRC) were excluded. Adjusted p-values < 0.5 are shown (Wilcoxon rank-sum test with Benjamini-Hochberg adjustment, excluding samples without *bai* operon detected). **f,** Relative abundances of bacterial species in colonic content of mice colonized with the BACOMI consortium with (7αDH+, n = 10, 5 female (F) and 5 male (M) mice) or without (7αDH-, n = 10, 5F/5M) *C. scindens* as measured by targeted qPCR. **g,** Primary (grey) and secondary (brown) BA profile in caecal content quantified using LC-MS/MS (female and male mice are shown separately; the stacked bars indicate mean values; all individual concentrations are available in **Table S4**). **h/i**, Expression of genes related to (**h**) BA metabolism and (**i**) epithelial cell proliferation and differentiation in the colonic mucosa. **j,** Exemplary images of immunofluorescence staining for Ki67+ cells in colonic crypts of BACOMI(7αDH-) or BACOMI(7αDH+) mice and corresponding quantification shown as bar plots (n = 4 mice each). **f, h, i, j** show mean values + standard deviation. Statistics: Wilcoxon rank sum test with Benjamini-Hochberg adjustment (* p.adj. ≤ 0.05; ** p.adj. ≤ 0.01, *** p.adj. ≤ 0.001, **** p.adj. ≤ 0.0001).

Due to the metagenomic data aforementioned, we used *C. scindens* in a gnotobiotic mouse model to study the relevance of DCA production in CRC development. Germfree wildtype mice on a high-fat diet were colonized with a simplified consortium of human gut bacteria referred to as the “BA-converting synthetic microbiota” (BACOMI) (**Fig. S1d**). BACOMI includes 7α-dehydroxylation activity (7αDH+) by *C. scindens* to produce DCA and can be compared to the condition without DCA when *C. scindens* is excluded from the consortium (7αDH-). Targeted qPCR analysis revealed a high relative abundance of 23.0 ± 2.9% (mean ± SD) for *C. scindens* in BACOMI-colonized mice and a relatively uniform colonization pattern of the other bacteria after 3 months (**Fig. 3f**). Consistently, the presence of *C. scindens* caused high levels of secondary BAs, in particular DCA, which was not detected when 7αDH activity was lacking in the microbial consortium (**Fig. 3g**). DCA concentrations were higher in female than male mice when colonized by *C. scindens*. Interestingly, 7αDH activity and DCA enhanced the expression of genes within BA-dependent signaling pathways (**Fig. 3h**) and genes known to be involved in tumorigenesis in the colonic epithelium (**Fig. 3i**). Furthermore, 7αDH activity was linked to higher numbers of Ki67+ epithelial cells in the colon (**Fig. 3j**).

### Microbially produced deoxycholic acid promotes tumorigenesis in the colon

To analyze whether 7αDH-mediated BAs affect colonic tumorigenesis, germfree wildtype mice on high-fat diet were colonized with BACOMI with or without *C. scindens* and then treated with AOM and dextran sodium sulphate (DSS) (**Fig. 4a**). To consolidate these findings independent of the 7αDH+ bacterial species used, we performed additional experiments in mice colonized with the mouse gut isolate *Extibacter muris* (Afrizal et al., 2022; Streidl et al., 2019) instead of *C. scindens*. The *bai* operon of *E. muris* has a high similarity to the *bai* operon of *C. hylemonae* (Streidl et al., 2021) and the latter species was also significantly associated with CRC in our metagenomic analysis (**Fig. 3c**). Moreover, *C. hylemonae* is proposed to be reclassified within the genus *Extibacter* in GTDB (Parks et al., 2022), highlighting the phylogenetic relatedness of these two species.

**Fig. 4.**
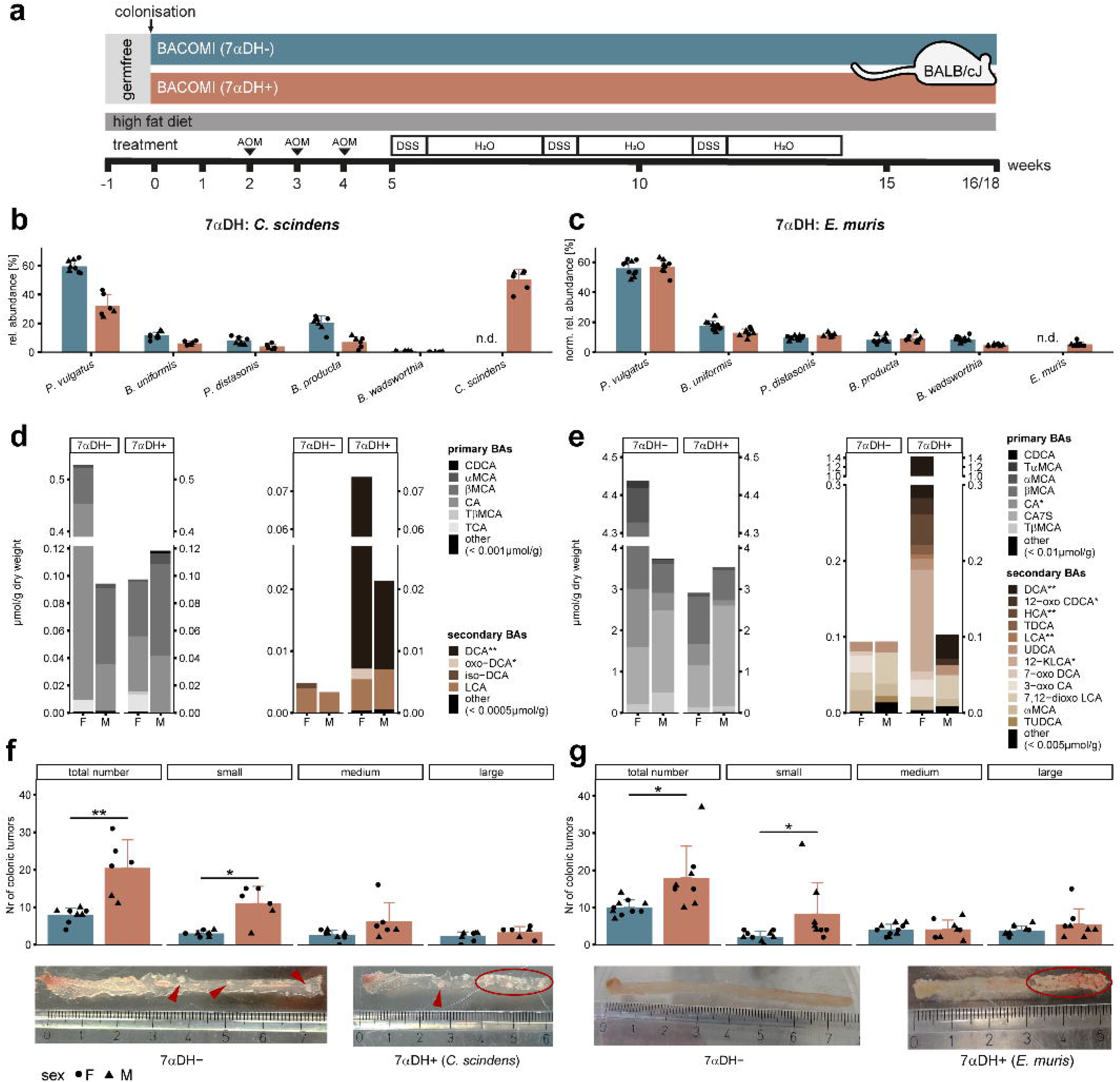
Effects of bacterial 7αDH on colonic tumorigenesis in gnotobiotic mice treated with AOM/DSS. **a**, Setup of AOM/DSS experiments using BACOMI(7αDH-) mice deprived of 7αDH activity and their BACOMI(7αDH+) counterparts colonised with either *C. scindens* (**b, d, f**, 7αDH-, n = 8-7; 7αDH+, n = 6) or *E. muris* (**c, e, g,** 7αDH-, n =10; 7αDH+, n = 8). Mice were culled after either 16 weeks (*C. scindens*) or 18 weeks (*E. muris*). **b, c,** Relative abundances of the bacterial species in colonic content as analysed by qPCR (**b**) or 16S rRNA gene amplicon analysis (**c**). **d, e,** Primary (grey) and secondary (brown) BA profile in caecal content as quantified using LC-MS/MS (female and male mice shown separately). Data are shown as stacked bar plots of mean values. Individual concentrations are available in **Table S4**. **f**, **g,** Number of colonic tumors; total or stratified according to their size: small (< 1.2 mm^2^), medium (1.2 - 2.5 mm^2^), large (> 2.5 mm^2^). Exemplary macroscopic pictures of the colon are shown below the bar plots; single tumors or areas with many tumors are indicated with red arrowheads or circles. In **b**, **c**, **f**, **g**, the sex of mice is indicated by different symbols (triangle = male; dots = female). Statistical analysis (**d-g**): Wilcoxon rank sum test with Benjamini-Hochberg adjustment (* p.adj. ≤ 0.05; ** p.adj. ≤ 0.01, *** p.adj. ≤ 0.001).

Following AOM/DSS treatment, *C. scindens* was present at higher relative abundance (50.3 ± 7.1%) (**Fig. 4b**) when compared with untreated BACOMI-colonized mice (**Fig. 3f**), suggesting favorable growth conditions for *C. scindens* under chemically induced colonic tumorigenesis. DCA was only detected in gnotobiotic mice colonized with *C. scindens*, including higher levels in female mice (**Fig. 4d**). Levels of oxo-DCA were also higher in the caecal content of female mice when *C. scindens* was present. Despite these sex differences in secondary BA concentrations, bacterial 7αDH activity led to higher tumor numbers in the colon of nearly all mice (**Fig. 4f**). Stratification according to tumor size revealed significance for small tumors, which suggests early-stage promotion of tumorigenesis rather than tumor progression.

Compared with *C. scindens*-colonized mice*, E. muris* was less abundant, which was linked to a more uniform distribution of the remaining bacterial species (**Fig. 4c**). DCA was detected in caecum when 7αDH activity by *E. muris* was present and sex-specific differences (higher levels in females) were confirmed (**Fig. 4e**). Several other secondary BAs were significantly elevated when *E. muris* 7αDH was present (e.g., 12-oxo CDCA, HCA, TDCA or LCA). Interestingly, levels of the primary BA CA were higher in 7αDH-mice, indicating enhanced conversion in 7αDH+ mice. Similar to the effects of *C. scindens*, 7αDH activity by *E. muris* caused higher numbers of tumors in the colon, which was again linked primarily to small tumors (**Fig. 4g**).

To confirm the role of microbial DCA production in CRC development, we performed gnotobiotic experiments in a further colonic tumor model using tissue specific nAtf6 transgenic mice (nAtf6^IEC^) (Coleman et al., 2018) with interleukin-10 knockout (*Il10*^-/-^) and a different synthetic community of cultured gut bacteria. Germfree IEC-specific Atf6 transgenic mice deficient for IL-10 (nAtf6^IEC^ ^tg/wt^;*Il10*^-/-^) colonized with *E. muris* in addition to the defined murine bacterial community OMM12 (Brugiroux et al., 2016) (OMM12+E) were characterized by more and earlier dropouts (**Fig. 5a**), higher fraction of responders (mice that developed tumors compared with non-responders) (**Fig. 5b**), and significantly higher numbers of tumors (**Fig. 5c** and **5d**). Interestingly, nAtf6^IEC^ ^tg/wt^;*Il10*^-/-^ mice colonized only with OMM12 and nAtf6^IEC^ ^fl/fl^;*Il10*^-/-^ colonized with *E. muris* (OMM12+E) had lower number of tumors, indicating that both, DCA-producing bacteria and gene susceptibility are required for colonic tumorigenesis.

**Fig. 5.**
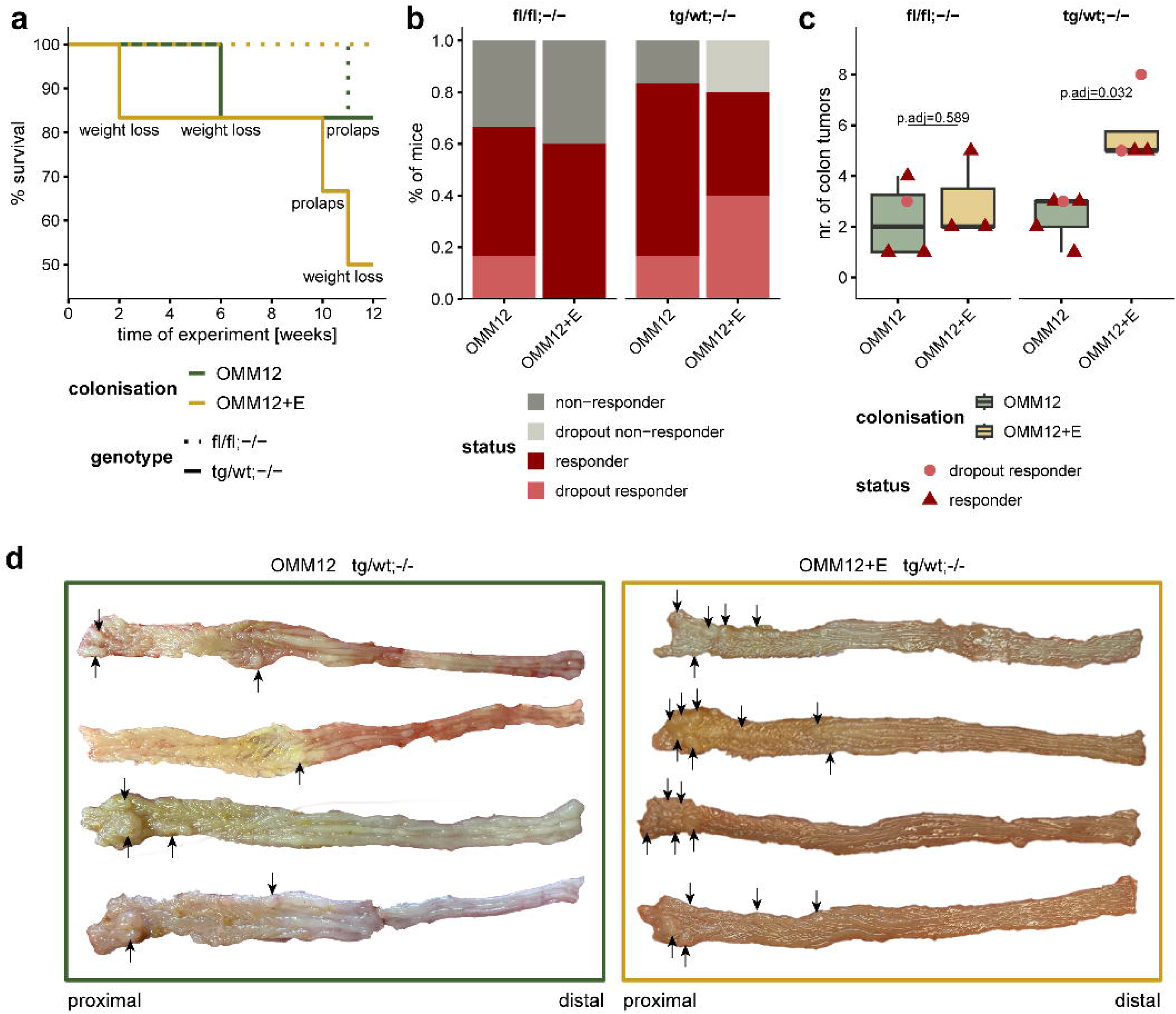
Effects of bacterial 7αDH activity on colonic tumor development in gnotobiotic model of colitis-associated cancer. Germfree nAtf6^IEC^ ^fl/fl^*;Il10^-/-^* (fl/fl;-/-) and nAtf6^IEC^ ^tg/wt^;*Il10*^-/--^ (tg/wt;-/-) mice were colonized either with the synthetic community OMM12 (n=6 per genotype) or OMM12+E (n=5 per genotype). **a** Survival curves of mice during the experiment. **b** Percentage of responders (mice with tumors) or non-responders (without tumors), including the respective fraction of early dropouts. **c** Number of colon tumors in responder mice. **d** Representative macroscopic images of the entire colon cut open longitudinally. Pictures of only tg/wt,-/- mice are shown; left, colonized with OMM12; right, colonized with OMM12+E. Tumors are indicated with black arrows. Statistics: Wilcoxon rank sum tests with Benjamini-Hochberg correction.

## Discussion

This study demonstrates the role of secondary BAs produced via gut bacterial 7αDH as tumor-promoting metabolites in experimental CRC. In contrast to non-physiological DCA supplementation in previous studies, we provide causal evidence linking the CRC risk-promoting effects of Western diets to functional adaptation of the gut microbiota. The experiments in *APC*^1311/+^ pigs substantiate the clinical relevance of Western diet-mediated stimulation of microbial BA metabolism and suggest that promoting the excretion of BAs is a therapeutic option to limit their tumorigenic effects in the large intestine.

Bacterial 7αDH activity from the specific DCA-producers *C. scindens* and *E. muris* enhanced colitis-associated CRC in gnotobiotic mice. This validates our own findings about the higher occurrence of *bai* operons from *C. scindens* and *C. hylemonae* in metagenomes from CRC patients and it is an important experimental proof of previous observational data on higher numbers of 7αDH+ bacteria in feces of CRC patients or individuals at high CRC risk (Hill et al., 1975; Kawano et al., 2010; Ocvirk et al., 2020; Ocvirk & O’Keefe, 2021; Ou et al., 2013; Wirbel et al., 2019). It also provides physiologically relevant evidence for the tumor-promoting effects of secondary BAs in murine CRC models compared with previously tested oral DCA supplementation and observed effects in the small intestine (Bernstein et al., 2011; Cao et al., 2014, 2017; Flynn et al., 2007). Of note, 7αDH caused significantly higher numbers of small but not large colonic tumors in our gnotobiotic mouse experiments with AOM/DSS treatment. This may indicate that DCA promotes cellular processes underlying tumor initiation rather than progression in the colonic epithelium of susceptible hosts. Data obtained in the nAtf6^IEC^ ^tg/wt^;*Il10*^-/-^ gnotobiotic model of CRC demonstrates the importance of host genetic modification besides gut inflammation for the tumor-promoting effect of 7αDH+ bacteria.

There are multiple possible mechanisms for the tumor-promoting effect of DCA. For instance, it was shown to stimulate cancer stemness in colonic epithelial cells by modulating β-catenin signaling (Farhana et al., 2016; Pai et al., 2004). It also triggered proliferation and DNA damage in *Apc*^Min/+^ Lgr5+ intestinal stem cells by antagonizing the BA receptor Fxr (T. Fu et al., 2019). Moreover, oral administration of DCA and high-fat diet-associated secondary BAs promoted proliferation of Lgr5+ stem cells via the BA receptor Tgr5 (J.-Y. Li et al., 2022). Consistently, 7αDH-mediated production of DCA did not induce colonic tumors without AOM/DSS in our study but it increased Ki67+ epithelial cell numbers and enhanced the expression of genes previously shown to be involved in secondary BA-associated colonic tumorigenesis (T. Fu et al., 2019; Pai et al., 2004). The tumor-promoting role of secondary BAs in susceptible hosts due to diet is also supported by our observation of Western diet-driven increase in Ki67+ cells in the colon of *APC*^1311/+^ pigs. Furthermore, enhanced fecal excretion of BAs by colestyramine was associated with reduced proliferation in the colonic epithelium. This finding provides additional pre-clinical evidence for the potential use of colestyramine for CRC prevention. Previous data in mice showed that colestyramine decreased the expression of genes involved in epithelial cell proliferation in the colon and that it could prevent streptozotocin and high-fat diet-induced hepatocellular carcinoma (J.-Y. Li et al., 2022; Xie et al., 2016). The use of BA sequestrants in individuals at increased CRC risk appears to be promising for prevention, since pre-diagnostic high plasma levels of BAs were linked to CRC outcome in a prospective study (Kühn et al., 2020). In this context of preventive BA reduction, it is worth noticing that secondary BAs also increased the expression of genes with tumor-suppressive or essential regulatory functions (e.g., *Guca2a*, *Muc2*, *Pparg*) in our gnotobiotic mice, suggesting that 7αDH-associated microbial metabolites have a fundamental role in regulating intestinal epithelium homeostasis and regeneration. Thus, low but not completely abolished DCA levels seem to be a promising preventive or therapeutic target for CRC risk reduction. Recently, BAs were shown to be required for TGR5-dependent regeneration of intestinal epithelial cells and protection from colitis in mice (Sorrentino et al., 2020), further supporting the importance of physiological BA levels.

In the mouse experiments, we detected a larger caecal BA pool and higher DCA concentrations in female compared to male animals. It is known that female and male mice differ in their BA composition and female mice tend to have higher serum and liver DCA concentrations (Z. D. Fu et al., 2012; Ma et al., 2020; Phelps et al., 2019; Selwyn et al., 2015). Moreover, sex hormones influence hepatic BA metabolism, including their production, secretion, and absorption (Z. D. Fu et al., 2012; Phelps et al., 2019). Therefore, our data is in line with already described sex differences in BAs and illustrates the importance to investigate both sexes. Besides BAs, sex hormones also affect CRC risk directly: testosterone increases CRC risk (Amos-Landgraf et al., 2014; C.-H. Song et al., 2021), whilst estrogen has the opposite effect (Chlebowski et al., 2004; Murphy et al., 2015; Son et al., 2019). This possibly explains why, in our gnotobiotic experiments, colonic tumor numbers were increased by secondary BA production in both male and female mice despite markedly higher DCA concentrations in the latter group.

Besides the causal role of secondary BAs in experimental CRC as demonstrated here, Western diet may promote CRC risk via the microbiome in several manners. For example, a recent study by Yang et al. (2022) (Yang et al., 2022) demonstrated that experimental CRC was associated with higher levels of *Alistipes* spp. and dysregulation of gut microbial co-metabolism leading to higher levels of glycerophospholipids. Moreover, diets high in fat content are usually associated with increased occurrence of *Enterobacteriaceae*, potentially including pks-expressing *Escherichia coli* with tumorigenic effects (Arthur et al., 2012). CRC risk does not exclusively arise from the increase in tumor-promoting factors but also from the lack of tumor-suppressive dietary compounds, including complex carbohydrates fermented by gut bacteria to SCFA such as butyrate (Donohoe et al., 2014; O’Keefe et al., 2015; Ocvirk et al., 2020). Furthermore, Western or high-fat diets are very diverse in terms of fat content and quality, with consequences regarding effects on the host. A recent study by Dmitrieva-Posocco et al. (2022) showed that a ketogenic diet, highly enriched in fat and very low in carbohydrates, reduced experimental colonic tumorigenesis due to anti-proliferative effects of the ketone body β-hydroxybutyrate. However, the long-term clinical relevance of ketogenic diets remains to be demonstrated. Taken together, these findings highlight the complexity of diet-associated changes in gut microbial metabolism and support the use of model organisms and experimental diets with detailed composition to unravel the influence of nutrition on CRC development or prevention.

In conclusion, our studies demonstrate the causal role of gut microbial 7αDH activity in experimental CRC. Secondary BAs, in particular DCA, are implicated in the tumor-promoting effects of Western diets, which provides a promising target for CRC prevention via dietary intervention or modulation of gut microbial BA metabolism.

## Methods

### Animal work

All experiments were carried out according to the German Animal Welfare Act and the European Union Normative for Care and Use of Experimental Animals. Individual approval numbers from the corresponding authorities are provided in the following subsections that describe the specific experiments.

### Porcine experiments

These experiments were approved by the Federal Government of Bavaria (permit no. ROB55.2-2-2532.Vet_02-18-33). Pigs were provided by the Chair of Livestock Biotechnology at the Technical University of Munich, Germany, and were housed and sampled at the animal facility Thalhausen (Faculty of Life Science Weihenstephan, Freising, Germany).

#### Dietary Interventions in *APC*^1311/+^ pigs

The experimental design is depicted in**. Fig. S1**. *APC*^1311/+^ pigs (German landrace × minipig crossbreed) (Flisikowski et al., 2022) underwent colonoscopy at the age of 3 months to enable their separation into two groups with a comparable distribution of individuals in terms of polyp numbers and size, sex, and littermates. Both groups were fed a standard diet until the dietary intervention started at the age of 4 months to allow for recovery after colonoscopy. The control (CTRL) and experimental diet enriched in red meat and lard (RL) were specifically designed for this experiment and purchased from ssniff Spezialdiäten GmbH (Soest, Germany; cat. no. S5745-S074 and S5745-S076, respectively). Main differences between the diets are illustrated in **Fig. S1** and their detailed compositions are provided in **Table S1.** The diets were fed to the pigs in a restricted manner (approx. 1 kg/d) for 3 months until the age of 7 months. At the end of the feeding period, fecal samples were taken, and colonoscopy was performed, including biopsy collection. Two RL pigs dropped out of the study due to tumor-independent reasons (meningitis and a foot malformation).

For the second feeding trial testing the effects of colestyramine, the overall experimental design was as in the first experiment (**Fig**. **S1**). However, *APC*^1311/+^ pigs were assigned to three groups instead of two (n = 4-5 pigs each): CTRL, RL, and COL (= RL diet supplemented with 12 g/kg colestyramine; Ratiopharm GmbH, Ulm, Germany; EAN 4150037520609). The recommended dose by the manufacturer is 4-16 g/day for adult subjects. Additional components of the drug colestyramine besides the active agent were added to CTRL and RL diet. Detailed diet compositions are available in **Table S1.**

#### Colonoscopy

Pigs underwent colonoscopy one month prior to dietary intervention and after the three months of experimental feeding. To reduce the load of fecal material in the large bowel, pigs were fasted for 24 h before the procedure. Based on bodyweight, pigs were narcotized (0.2 mg/kg ketamin and 0.05 mL/kg azaperon) and the large bowel was flushed with tap water to remove remaining luminal content. Tissue biopsies and images were taken using a STORZ colonoscopy system (Flexible SILVER SCOPE^®^).

#### Collection of fecal samples and sampling

In both pig trials, fresh fecal samples were collected prior to (4 months of age) and immediately after dietary intervention (7 months of age) before colonoscopy. In brief, fecal material was taken directly from the rectum of each pig using sterile gloves and placed into individual, sterile plastic containers. Solely the material from core areas of the feces was collected using a sterile spatula. Samples were immediately frozen on dry ice for later sequencing and metabolomics and stored at -80°C. Tissue samples from the rectum were collected either during colonoscopy (pig trial 1) or after the pigs were culled (pig trial 2). The resulting samples were formalin-fixed and paraffin embedded tissue blocks were stored at room temperature.

### Mouse experiments

#### BACOMI consortium

The synthetic microbial community BACOMI (Bile acid-converting microbiota) contains six bacterial species (all obtained from the Leibniz-Institute DSMZ - German Collection of Microorganisms and Cell Cultures, Germany) that can transform the host-derived primary BAs into dehydroxylated secondary BAs (adapted from Ridlon et al. (Ridlon et al., 2020)) (**Fig. S1d**). *Bacteroides uniformis* (*B. uniformis*) DSM 6597, *Phocaeicola vulgatus* (*P. vulgatus*) DSM 1447 and *Parabacteroides distasonis* (*P. distasonis*) DSM 20701 express bile salt hydrolases (BSH) that produce free primary BAs from conjugated counterparts. *Bilophila wadsworthia* (*B. wadsworthia*) DSM 11045 can further metabolize taurine released by BSH activity into H_2_S. *Clostridium scindens* (*C. scindens*) DSM 5676 is the key-member of the BACOMI consortium, as it is the only species with 7α-dehydroxylating (7αDH) activity. Finally, *Blautia producta* (*B. producta*) DSM 2950 was added due to being a dominant human gut bacterium that is H_2_-consuming, acetogenic, able to isomerize bile acids and thus increases variability of the BA pool. The other 7αDH+ species *Extibacter muris* (*E. muris*) DSM 28560 was isolated and described by us and available in the in-house collection (Lagkouvardos, Pukall, et al., 2016; Streidl et al., 2021). All bacteria except for *B. wadsworthia* were cultured in brain heart infusion (BHI) medium supplemented with resazurin, hemin, yeast extract and L-cysteine under anaerobic conditions. *B. wadsworthia* was cultured as previously described (Burkhardt et al., 2021).

#### OMM12 +/- *E. muris* consortium

OMM12 is a synthetic community of cultured bacteria representing the 5 major phyla in the mouse gut (Brugiroux et al., 2016). We amended this consortium by the addition of the 7αDH+ species *Extibacter muris* DSM 28560^T^ (*E. muris*). For the generation of cryo- stocks, individual strains were grown in anaerobic Akkermansia medium (Brugiroux et al., 2016). Strains were mixed equally (based on OD measurements) and mixed with 20% glycerol in a 1:1 ratio (exact mixing volumes and strain designations are given in **Table S7**).

#### Gnotobiotic mouse experiments

The experiments in AOM/DSS treated mice were approved by the Ministry of Social Affairs, Health, Integration and Consumer Protection of the state Brandenburg (permit no. 2347-15-2021). Germfree male and female wildtype mice (BALB/cJ) were kept in positive-pressure isolators at the animal facility of the German Institute for Human Nutrition Potsdam-Rehbruecke (Nuthetal, Germany) with a 12 h light-dark cycle at 22 ± 2 °C and 55 ± 5 % air humidity. Starting 1 week prior to colonization, the mice were switched from standard diet to a high-fat diet (HFD, **Table S1)**. Mice (n = 10 per group) were colonized by gavage (10^8^ cells per bacterium in 100 µL medium; 50 µL orally and 50 µL rectally at day 0 and 2) with BACOMI either with (7αDH+) or without (7αDH-) *C. scindens*. After colonization, mice were either kept for 13 weeks (**Fig. S1 and Fig. 5**) or subjected to further treatment to induce experimental CRC (**Fig. 6**). In the latter case, mice received three intraperitoneal injections of azoxymethane (AOM) (5 mg/kg body weight) in the interval of one week, starting at week 2 after colonization. Five days after the final AOM injection, mice were treated with 3 cycles of dextran sodium sulphate (DSS) with 5 days of DSS (1.5 % in drinking water) followed by 16 days of normal drinking water. Mice were culled two weeks after the end of the AOM/DSS treatment and colonic tumors were counted using a binocular microscope. The colonization and AOM/DSS treatment of mice with *E. muris* followed the same protocol, but mice were kept 2 weeks longer after colonization and prior to AOM/DSS treatment to confirm colonization status. For BACOMI with *C. scindens* colonization, three mice (male n = 1, female n = 2) dropped out in the 7αDH+ group during the first DSS cycle, whereas for the 7αDH-group, two mice (male n = 1, female n = 1) dropped out at the end of the third DSS cycle or after the first AOM injection, respectively. For BACOMI with *E. muris* colonization, two female mice dropped out in the 7αDH+ group after the first DSS cycle.

Germ-free IEC-specific expression of activated (p50) nAtf6 transgenic mice (nAtf6^IEC^) (Coleman et al., 2018) were crossed to germ-free *Il10*^-/-^ mice. Heterozygous mice with monoallelic nAtf6 expression (nAtf6^fl/wt^;Vil-Cre^tg/wt^;*Il10*^-/-^) and floxed controls (nAtf6^fl/wt^;Vil- Cre^wt/wt^;*Il10*^-/-^ ) were used for gnotobiotic work. These experiments were approved by the Committee on Animal Health Care and Use of the state of Upper Bavaria (Regierung von Oberbayern; AZ TVA 55.2-2532.Vet_02-18-121). Four-week-old mice were inoculated orally (50µL) and rectally (100µL) with cryo-stocks of the OMM12 or OMM12 + *E. muris* (OMM12+E) strain mixture at day 0 and day 3, and kept colonized for 12 weeks. All mice were euthanized with CO_2_ at the end of an experiment (age 16 weeks) or when abortion criteria were met.

### Quantitative real-time PCR for intestinal gene expression

The Dynabeads mRNA purification kit (ThermoFisher Scientific, USA) was used for mRNA isolation. Lysis buffer (200 µl) and 10 zirconium beads were added to approximately 5 mg of intestinal tissue stored at -80 °C in RNA later tissue reagent (Qiagen, Germany). A Tissue lyser (Qiagen, Germany) was then used for homogenization (2 × 2 min at 50 Hz with a 2 min break in between). The lysate was pulled through a syringe five times with a 1 mL syringe and centrifuged (14,000 rpm, 2 min, room temperature). The rest of the protocol was as per the manufacturer’s instructions. mRNA concentrations were determined using a Nanodrop (Peqlab VWR, USA) and 20 ng were used for reverse transcription using the RevertAid H Minus First Strand cDNA Synthesis kit (ThermoFisher Scientific, USA) according to manufacturer’s recommendations. For qPCR, the mastermix was prepared according to the manufacturer’s recommendations given by the Quantinova SYBR Green PCR Kit (Qiagen, Germany), and forward and reverse primers added (0.5 µM each) (**Table S5**). Then, 4 ng template cDNA was used in a total reaction volume of 10 µl. Samples were amplified using the 7500 Fast Real-Time PCR System (Applied Biosystems, USA). To determine C_T_-values, the Life Technologies 7500 Software (version 2.3) was used. Normalization was to the housekeeping gene *Rpl13a,* which encodes for the 60S ribosomal protein L13a. Results were calculated relative to the control group of 7αDH- mice.

### Immunohistochemistry (IHC) and immunofluorescence (IF)

Specimens from the two pig trials were fixed in 4 % formaldehyde solution for 24 h, embedded, and sectioned (3 µm). The tissue sections were then deparaffinized, antigen unmasked (citrate buffer, pH = 6) and endogenous peroxidases were inactivated in 3% H_2_O_2_ for 10 min. After blocking for 1 h (2% goat serum in PBS), sections were incubated overnight at 4 °C with the primary antibody for Ki67 (Rabbit, DSC innovative Diagnostik-System KI681C002, dilution 1:400) followed by a second overnight incubation with the secondary antibody (Goat Anti-Rabbit IgG, Santa Cruz sc-2780, dilution 1:400). Peroxidase activity was detected using diaminobenzidine (DAB) substrate or the VECTASTAIN^®^ Elite^®^ ABC Kit (Vectorlab PK-6100), respectively. Staining of Ki67 was considered positive and quantified only when detected in nuclei. Ten randomly selected fields (magnification x40) from each tissue section were analyzed. Values are presented as the percentage of the stained cells per area.

For the gnotobiotic mouse trial (**Fig. 5**), the dissected tissues were fixed using Methacarn solution and embedded in paraffin. Tissues were cut into 4 µm-thick sections and deparaffinized by heating (60 °C, 15 min) followed by two times xylol for 3 min. The specimens were then rehydrated using a decreasing alcohol series: 100 % for 2 min, 96 % for 2 min, and 70 % for 1 min, followed by water for 1 min. After rinsing twice in water (5 min each), the specimens were boiled in 10 mM citrate buffer (pH = 6) for 30 min for antigen retrieval, followed by 30 min of cooling. Thereafter, the specimens were washed 3 times with water for 5 min, once with PBS for 5 min, and then blocked with PBS containing 5 % goat serum for 1 h at room temperature. Primary antibodies against Ki67 (ab15580, dilution 1:200) were incubated overnight at 4°C. Specimens were then washed three times in PBS (5 min each) and incubated with the secondary antibody (AlexaFluor-488, ThermoFisher, dilution 1:200), and Hoechst (dilution 1:200) for 1-2 h at room temperature. All antibodies were diluted in PBS containing 1 % bovine serum albumin and 0.3 % Triton X-100. Finally, the specimens were washed in PBS for 5 min each and cover with glass slides using mounting medium (Vector Laboratories, United States). For imaging, a confocal laser microscope was used (LSM 780 microscope, Zeiss, Germany). Tile scans with 40x magnification were acquired and analysed with the ZEN (black edition) 2.3 software (Zeiss, Germany).

### Microbiota composition analysis

Metagenomic DNA was obtained using a modified version of a previously published protocol (Godon et al., 1997; Just et al., 2018). Cells were lysed mechanically via bead-beating and DNA was purified on columns (Macherey-Nagel, Germany). A robotized platform (Biomek4000, Beckman-Coulter, Germany) was then used for library construction. The V3-V4 regions of 16S rRNA genes were amplified (25 cycles) in a two-step PCR (Berry et al., 2011) using primers 341F (5′-cctacgggnggcwgcag) and 785R (5′-gactachvgggtatctaatcc) (Klindworth et al., 2013) and 24 ng of template DNA. A double combinatorial indexing strategy was used. Amplicons were purified using the AMPure XP system (Beckmann Coulter, Germany), pooled in equimolar amounts and sequenced in paired-end mode using a MiSeq system (Illumina).

Raw data were processed using IMNGS (https://www.imngs.org) (Lagkouvardos, Joseph, et al., 2016), an online platform based on UPARSE (Edgar, 2013). Sequences were clustered into operational taxonomic units (OTUs) at 97% sequence similarity and only those with a relative abundance ≥ 0.25% in at least one sample were kept to exclude spurious OTUs (Reitmeier et al., 2021). Parameters for data processing were: min/max sequence length, 300-500nt; barcode mismatch, 1; end-trim length, 15; max. no. expected errors, 3; trimming q score, 3. The number of sequences per sample after filtering was 26,286 ± 8,044 for the first pig trial and 12,564 ± 3,954 for the gnotobiotic mouse experiment with *E. muris*. Downstream diversity and composition analyses were performed in Rhea (https://github.com/Lagkouvardos/Rhea) (Lagkouvardos et al., 2017). For the first pig trial, taxonomies were assigned using SILVA release 128 (Pruesse et al., 2012). Several families showed a significantly different behavior between the feeding groups over the time. A pseudocount (lowest value × 10) was added to the normalized relative abundance of each family per sample before calculation to avoid division by zero. For family comparison over time, a start-adjusted delta-value was calculated as follows: (end – start) / start. For changes in α-diversity, original values without pseudo-count were used for the same calculation. Identities of the OTUs for the gnotobiotic mouse experiment were confirmed using Eztaxon (Yoon et al., 2017). Normalized relative abundances are plotted in **Fig. 2** and **Fig. 6** and provided in **Table S2** and **Table S3.**

For the gnotobiotic mouse trials with *C. scindens,* the QIAmp Fast DNA Stool Mini Kit (Qiagen, Germany) was used to extract and purify DNA from colon content. Approximately 50-80 mg frozen content was added to a cryo-tube containing ca. 100 mg of 0.1 mm glass beads, and 1 mL of InhibitEx buffer, followed by bead-beating (Uniprep, UniEquip, Germany) for 2 × 5 min at 3,000 rpm. All samples were processed according to manufacturer’s instructions except for the elution volume (changed to 70 µL). For DNA isolation from bacterial cultures, the Genomic DNA from Microorganisms Kit (Macherey-Nagel, Germany) was used as per the manufacturer’s instructions. The disruption time was 12 min, and the samples were eluted twice using the same elution buffer.

A standard curve plotting bacterial cell number against C_T_-value was established for each BACOMI species using RT-qPCR and species-specific primers (**Table S5**). The standard curve was generated from overnight bacterial cultures and cell numbers were determined using a Thoma cell counting chamber (Fein-Optik, Germany). For quantification, bacterial DNA was isolated from colon content and bacterial cell numbers (**Table S4**) were calculated using the generated standard curve.

### Analysis of *bai* operon occurrence in metagenomes

To quantify the occurrence of *bai* operons in fecal metagenomes, we first extracted *bai* sequences from Kim et al., (Kim et al., 2022) and retained the operons harboring at least 7 of the 8 genes found in experimentally validated *bai* operons. The complete contig of each *bai* sequence was downloaded from GenBank. GECCO (v0.9.7) was trained on the complete contigs with default parameters using Pfam v35.0 (Mistry et al., 2021) and *bai*-specific HMMs (Wirbel et al., 2019) as training features. We then ran GECCO on ProGenomes3 (PG3) representative genomes (n = 41,171) and selected those clusters with more than seven unique *bai* genes, since all experimentally verified *bai* operons had >7 unique *bai* genes with our approach, based on *bai* genes HMM annotation. This resulted in a total of 18 PG3 genomes with complete *bai* operons and we extracted the protein sequence of each of the individual operons. We then computed protein sequence similarity across the *bai* genes between operons from all genomes to understand how divergent the sequences are, which is important to know prior to quantifying relative abundances in metagenomes. Protein sequence similarity was computed by pairwise alignment (global mode, BLOSUM62 substitution matrix) as implemented in Biopython v1.81 (Cock et al., 2009). Next, we used mmseqs map (v13.45111) to map protein sequences of the complete operons of all 18 clusters against our reduced GMGC (Global Microbial Gene Catalog) gut catalogue (n = 13,788,251 ORFs) to identify which genes contained parts of the operons. This approach also enabled species-resolved *bai* operon quantification. We then collected a total of n = 2,142 samples from case-control metagenomic studies of CRC (n = 1,034 CRC, n = 1,108 CTRL, see **Table S8** for an overview). These metagenomes were processed and profiled as follows: (i) Raw reads were cleaned using bbduk (v38.93), including low-quality trimming on either side (qtrim=rl trimq=3), discarding low-quality reads (maq=25), adapter removal (ktrim=r k=23 mink=11 hdist =1 tpe=true tbo=true; against the bbduk adapter library), and length filtering (ml=45); (ii) Reads were screened for host contamination using kraken2 (v2.1.2) against the human hg38 reference genome with ribosomal sequences masked (Silva_v_138); (iii) Cleaned reads were then mapped onto our reduced human gut gene catalogue using BWA-MEM (v0.7.17) with default parameters and alignments were filtered to >45bp alignment length and >97% sequence identity. Reads aligning to multiple genes contributed fractional counts towards each hit gene. Alignment counts for a gene were normalized against its length, then scaled up according to the strategy employed by NGLess (https://ngless.embl.de/Functions.html#count) and propagated to the functional features with which the gene is annotated. Extensive use of gffquant (v.2.10.0, https://github.com/cschu/gff_quantifier) was made throughout the functional profiling workflow. As information about which individual GMGC gene corresponded to a part of the *bai* operon of a given bacterial species was available, we obtained species-resolved relative abundances of *bai* operons. Values for all GMGC genes belonging to a specific species’ *bai* operon were summed to compute total relative abundance for the *bai* operon of the given bacterial species. We only included *bai*-encoding species if they occurrence at a prevalence of at least 2% in the entire dataset. To calculate prevalence of the bacterial species, we performed taxonomic profiling using mOTUs (v3.1) with default parameters after having run the bbduk quality filtering as described above. Differential abundance analysis was performed using a linear mixed model with the study as a random effect, through the SIAMCAT package. A pseudocount of 1e-8 was used for plotting zero values on a logarithmic scale.

### BA measurements

Pig stool samples from trial 1 and 2 were analyzed as described previously by Wegner et al. (2017) and Reiter et al. (2021), respectively (Reiter et al., 2021; Wegner et al., 2017).

For the BACOMI mouse experiments with *C. scindens,* lyophilized caecal content (125 mg) were mixed with 450 µL acetonitrile, followed by 15 min vortexing and then centrifugation (13,000 × g, 60 min, 21 °C). Supernatant (300 µL) was mixed with the internal standards (ISTDs) d_4_-CDCA and d_4_-LCA (10 µM each) and dried in a SpeedVac system (Jouan RC10.22, ThermoFisher Scientific, USA). The residue was dissolved with methanol and water [1:1]. Chromatographic separation of 5 µL sample was achieved on a 1290 Infinity II HPLC (Agilent Technologies, Germany) equipped with a Poroshell EC-C18 column (Agilent Technologies) (3.0 × 150 mm, 2.7 µm) connected to a guard column (3.0 × 5 mm, 2.7 µm) of the same material. The column was tempered to 30 °C. A mobile phase system consisting of 10 mM ammonium acetate/acetonitrile (80:20 v:v; solvent A) and water/acetonitrile (20:80 v:v; solvent B), was applied at a flow rate of 0.2LmL/min. The initial solvent composition of 85% solvent A was maintained for 6 min. Then, solvent A was decreased stepwise to 70, 40 and 20% within 20, 30 and 40 min of total run-time, respectively. After 45 min, the mobile phase reached initial conditions again. MS/MS analysis was carried out using a 6495C triple-quadrupole mass spectrometer (Agilent Technologies) operating in the negative electrospray ionization mode (ESI-). The ion source parameters were: sheath gas temperature, 400 °C; sheath gas flow, 12LL/min of nitrogen; nebuliser pressure, 40Lpsi; drying gas temperature, 120L°C; drying gas flow, 15LL/min of nitrogen; capillary voltage, 4500 V; nozzle voltage, 0 V; iFunnel high pressure RF voltage, 90 V and iFunnel low pressure RF voltage, 60 V. A total of 20 compounds (18 bile acids and 2 ISTDs) were simultaneously analyzed by single reaction monitoring (SRM) or, if feasible, by multiple reaction monitoring (MRM). HPLC-MS/MS parameters are given in **Table S6.** Quantification was performed using MassHunter Workstation Quantitative Analysis for QQQ (Version 10.1, Agilent Technologies). Analytes were externally calibrated in the concentration range of 0.09 to 90 µM (ISTDs constantly 10 µM). BA concentrations were normalized to dry weight of the caecal content subjected to extraction.

For the gnotobiotic mouse experiment with *E. muris, s*amples were lyophilized overnight at –60 °C. The dried caecal contents were weighed, and 6 ceramic beads (2.5 mm) were added to each tube. Proportionally to the weight of each sample, between 500 µL and 1500 µL of MeOH/H_2_O (2/1) + 0.1% formic acid was used as extraction solvent. Samples were homogenized in a Precellys 24 Tissue Homogenizer (Bertin Instruments, France) at 6500 rpm 2 x 20” beat and 20” rest. The homogenized caecal samples were centrifuged at 21,000 rcf, for 15 min, at 4 °C. 100uL from each supernatant or calibration standard were transferred into individual wells of 2 mL 96-well plate. 50 µL of an ISTD solution (CA-d_4_, CDCA-d_4_, TCA-d_4_, TUDCA-d_4_, DCA-d_4_ and LCA-d_4_, each at 2 µM in methanol) was pipetted in each well. Immediately after the addition of ISTD, 600 µL of 0.2% formic acid in H_2_O was added to each sample or calibration standard level. The 96-well plate was shaken with an orbital shaker at 300 rpm and centrifuged at 3500 rpm, 5 min, 4 °C. The contents of the 96-well plate were extracted by solid phase extraction with an Oasis HLB 96-well uElution plate (Waters, USA). The extracted samples were dried in a Biotage® SPE Dry 96 (Biotage, Sweden) at 20 °C and reconstituted with 100 µL of MeOH/H_2_O (50/50). The plate was shaken with an orbital shaker at 300 rpm, 5 min and centrifuged at 3500 rpm, 5 min, 4 °C. The samples were injected on the LC-HRMS system.

The quantitative method was performed on an Agilent ultrahigh-performance liquid chromatography 1290 series coupled in tandem to an Agilent 6530 Accurate-Mass Q-TOF mass spectrometer (Vico-Oton et al., 2022). The separation was done on a Zorbax Eclipse Plus C18 column (2.1 × 100 mm, 1.8 µm) and a guard column Zorbax Eclipse Plus C18 (2.1 × 5 mm, 1.8 μm), both provided by Agilent technologies (USA). The column compartment was kept heated at 50 °C. Two different solutions were used as eluents: ammonium acetate [5 mM] in water as mobile phase A and pure acetonitrile as mobile phase B. A constant flow of 0.4 mL/min was maintained over 26 min of run time with the following gradient (expressed in eluent B percentage): 0-5.5 min, constant 21.5% B; 5.5-6 min, 21.5-24.5% B; 6-10 min, 24.5-25% B; 10-10.5 min, 25-29% B; 10.5-14.5 min, isocratic 29% B; 14.5-15 min, 29-40% B; 15-18 min, 40-45% B; 18-20.5 min, 45-95% B; 20.5-23 min, constant 95% B; 23-23.1 min, 95-21.5% B; 23.10-26 min, isocratic 21.50% B. The system equilibration was implemented at the end of the gradient for 3 min in initial conditions. The autosampler temperature was maintained at 10 °C and the injection volume was 5 µL. The ionization mode was operated in negative mode for the detection using the Dual AJS Jet stream ESI Assembly. The QTOF acquisition settings were configured in 4 GHz high-resolution mode (resolution 17000 FWHM at m/z 1000), data storage in profile mode and the high-resolution full MS chromatograms were acquired over the range of m/z 100-1700 at a rate of 3 spectra/s. The mass spectrometer was calibrated in negative mode using ESI-L solution from Agilent technologies every 6 h to maintain the best possible mass accuracy. Source parameters were setup as follows: drying gas flow, 8 L/min; gas temperature, 300 °C; nebuliser pressure, 35 psi; capillary voltage, 3500 V; nozzle voltage, 1000 V. Data were processed afterwards using the MassHunter Quantitative software and MassHunter Qualitative software (Agilent technologies, USA) to control the mass accuracy for each run. In the quantitative method, 43 bile acids were quantified by calibration curves. The quantification was corrected by addition of internal standards in all samples and calibration levels. Extracted ion chromatograms were generated using a retention time window of ± 1.5 min and a mass extraction window of ± 60ppm around the theoretical mass of the targeted bile acid.

For all experiments, BAs that were not detected by the respective methods were assumed to absent, set to 0 µmol/g and only those BAs detected in at least 50 % of the animals in at least one group were considered for downstream analyses. Statistics were done only on dominant BA species (the concentration thresholds are shown in the figures; all individual values for all BAs are available in **Table S2-4**). UDCA is a primary BA in rodents, but a secondary BA in humans (Honda et al., 2020; J. Li & Dawson, 2019). Due to their similarity to humans, we classified UDCA as a secondary BA in pigs.

### Lipidomic analyses

Fecal homogenates for analysis of fatty acids, sterols and stanols were prepared in isopropanol as described before (Schött et al., 2018). Subsequently, the homogenates were used for both fatty acid and sterol/stanol analysis. Concentrations of total fecal fatty acids were determined by gas chromatography coupled to mass spectrometry (GC-MS) as described previously (Ecker et al., 2012) with some modifications. Samples were derivatized to fatty acid methyl ester (FAME). The initial column temperature of 50 °C was held for 0.75 min, increased with 40 °C/min to 110 °C, with 6 °C/min to 210 °C, with 15 °C/min to 250 °C and held for 2 min. Iso- and anteiso-FAMEs standards were applied to identify branched chain fatty acids and to calibrate the instrument response. Samples for analysis of sterols and stanols were analyzed as described previously (Kunz & Matysik, 2019).

### Data visualization and statistics

Statistical analyses were done using R (Version 4.2.1) (R Core Team, 2022) in RStudio (Posit team, 2022) using the rstatix package. For microbiota analyses, detailed information is provided in Rhea (https://github.com/Lagkouvardos/Rhea) (Lagkouvardos et al., 2017). Comparisons between two groups were done by Wilcoxon tests and Benjamini-Hochberg adjustment. Comparisons between three groups were done by Kruskal Wallis tests with Dunn’s multiple comparisons and Benjamini-Hochberg adjustments. Adjusted p-values are with stars as defined in the corresponding legends; non-adjusted p-values are given as numbers directly in the graphs whenever appropriate. All graphs were created in R using tidyverse (ggplot2; RRID:SCR_014601) (Wickham H, 2016), ggpubr, cowplot, and ggbreak (Xu et al., 2021).

### Data availability

All data used to generate the figures is available in the supplementary tables (**Table S2-4**). Raw data of the 16S rRNA gene amplicon sequencing were submitted to ENA and are available under project PRJEB60380.

## Acknowledgements

We are grateful to Viola and Steffen Loebnitz, Tatiana Flisikowska, and Thomas Winogrodzky from the Chair of Livestock Biotechnology at the Technical University of Munich, Germany, and Dieter Saur from the Klinikum rechts der Isar, Technical University of Munich, Germany, for their support with pig experiments. We also thank Ines Grüner, Anika Sander, Sabine Schmidt and Anke Gühler from the German Institute of Human Nutrition Potsdam-Rehbruecke, Nuthetal, Germany, for their help with the germfree mouse colony and colonization experiments, and Tobias Goris and Annett Braune for their critical review of data.

## Funding

TC received funding from the German Research Foundation (DFG): project no. 460129525, NFDI4Microbiota; project no. 395357507 - SFB1371 “Microbiome Signatures”; and project no. CL481/8-1. TC, GZ, DH, and KF received funding from the German Ministry for Research and Education (BMBF): consortium project Mi-EOCRC (KZ. 01KD2102D). SO and DH received funding from the DFG: project no. 338582098 and no. 395357507 - SFB1371 “Microbiome Signatures”, respectively.

## Notes

### Competing Interest Statement

The authors have declared no competing interest.

### Summary of Updates

Additional data/experiments were added.

