## Supplementary material for "Secondary bile acid production by gut bacteria promotes Western diet-associated colorectal cancer": Fig. S1

### Supplemental Information

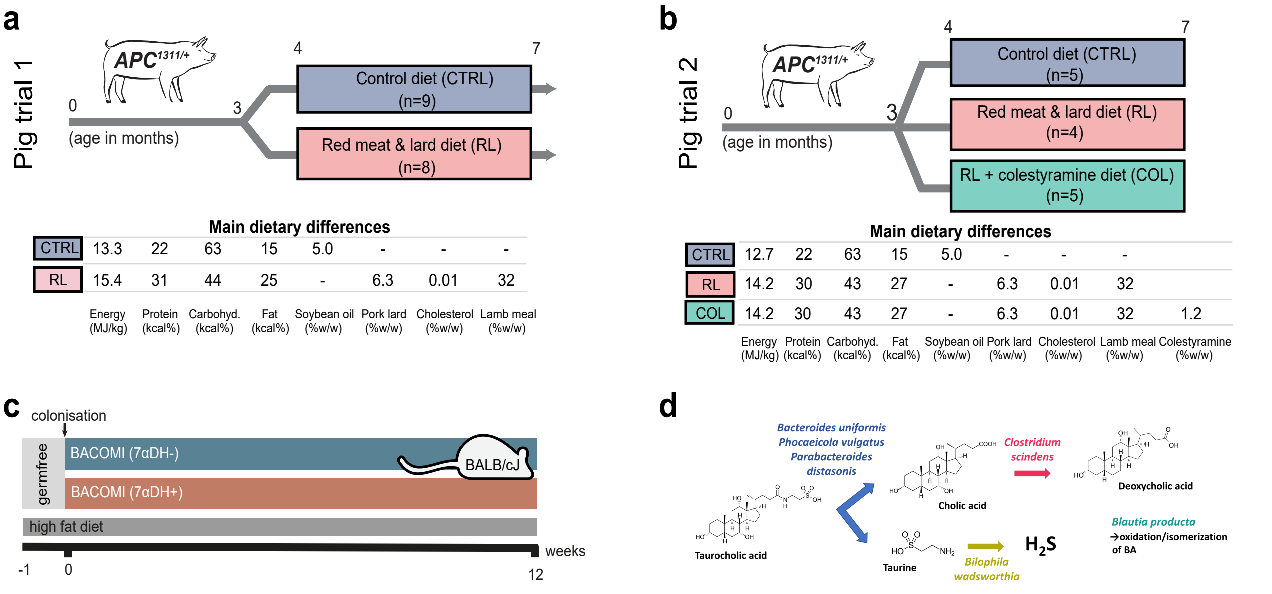


Panel a, Main dietary differences:

| | Energy (MJ/kg) | Protein (kcal%) | Carbohyd. (kcal%) | Fat (kcal%) | Soybean oil (%w/w) | Pork lard (%w/w) | Cholesterol (%w/w) | Lamb meal (%w/w) |
|---|---|---|---|---|---|---|---|---|
| CTRL | 13.3 | 22 | 63 | 15 | 5.0 | - | - | - |
| RL | 15.4 | 31 | 44 | 25 | - | 6.3 | 0.01 | 32 |

Panel b, Main dietary differences:

| | Energy (MJ/kg) | Protein (kcal%) | Carbohyd. (kcal%) | Fat (kcal%) | Soybean oil (%w/w) | Pork lard (%w/w) | Cholesterol (%w/w) | Lamb meal (%w/w) | Colestyramine (%w/w) |
|---|---|---|---|---|---|---|---|---|---|
| CTRL | 12.7 | 22 | 63 | 15 | 5.0 | - | - | - | |
| RL | 14.2 | 30 | 43 | 27 | - | 6.3 | 0.01 | 32 | |
| COL | 14.2 | 30 | 43 | 27 | - | 6.3 | 0.01 | 32 | 1.2 |

**Fig. S1** Experimental Design of **a**, pig trial 1, **b**, pig trial 2, including main differenes between the diets. **c**, Experimental design of the BACOMI trial without AOM/DSS treatment. **d**, Functional relevance of each BACOMI strain (adapted from Ridlon et al., 2020).
